# Trends in forest carbon offset markets in United States

**DOI:** 10.1101/2022.07.21.500541

**Authors:** Lilli Kaarakka, Julia Rothey, Laura E. Dee

## Abstract

Natural climate solutions are gaining international policy attention – with forests highlighted as a primary pathway for storing carbon. However, evaluations of additional carbon benefits and the permanence of forest carbon offsets projects remain scarce. In response, we compiled a novel database to analyze trends in existing forest management projects from the two largest offset project registries in the only carbon market in United States. We find that improved forest management projects represent 96% of all credits from forestry projects and 58% of all credits and span diverse practices with different potential for carbon storage. Our results also show that 26% of existing forest C offsets in the US are at risk from wildfire. From a policy perspective, our results underscore the need for more sophisticated insurance mechanisms for forest carbon offset reversals, and for a framework to monitor and evaluate cumulative and future carbon benefits of forest-based offset projects.

**Significance:** We assess trends in ownership, forest management practices and disturbance risks in existing forest carbon offset projects in the US.

## INTRODUCTION

Nature-based solutions have recently gained attention in science and policy arenas as a way to offset carbon (C) emissions arising from other industries ^1,2^. The demand for C offsets from nature-based solutions is expected to continue to increase as buyers prefer projects that demonstrate co-benefits beyond emission reductions (e.g., other ecosystem services beyond C) ^3–5^. In particular, carbon offset credits from forests have been in the spotlight. They are currently the primary type of projects in the global offset markets and have increased globally by 159% between 2020 and 2021 ^6^. For example, forest-based offsets represent 92% of offset credits issued in the Cap-and-Trade Program in California, USA. Given this rise in investment, an assessment of the trends and knowledge needs for these growing C offsets markets for forest projects is urgently needed.

Forest carbon offset credits are being issued for projects store *additional* carbon relative to the status quo, including for avoided forest conversion, reforestation, and improved forest management. Improved forest management – in the market-context broadly defined as any forest management activity that increases C stocks on forested land – is the most common forest C offset project type in the US ^7,8^. Improved forest management is an umbrella term for many forms of forest management and silvicultural practices, ranging from thinning to selection harvesting (reviewed in Kaarakka et al., 2021). These practices differ in their ability to store C relative to business-as-usual forest management and regulations in a region ^9,10^. However, evidence on the extent to which different improved forest management practices provide *additional* C benefits remains patchy (Kaarakka et al., 2021). Further, most prior analyses of nature-climate solutions from forest management only consider one form of management (extended rotations) ^2,11^, but a broader array of practices are being implemented or considered on-the-ground (Table 1). As a result, the types of improved forest management projects that have been credited for offsets (see Table 1) and their implications for C additionally remain to be quantified and verified.

**Table 1.** Forest management terminology used in project documentation for existing forest C offset projects, and the total forest carbon offsets credits issued (as of December 2020) per management practice/activity mentioned in project documentation. Note that some of the offset project documents mention multiple management practices and/or activities, so some offsets are listed multiple times in the table.

| Management practice or activity | Definition | Projects mentioned one or more of these forest management terms | Total offsets issued | % of offset credits issued |
| --- | --- | --- | --- | --- |
| No management or no commercial harvest | No forest management or commercial harvest is applied. | No management, no commercial harvest, no harvest | 62,721,277 | 34% |
| Uneven-aged management | Stands that have three or more age classes throughout the cutting cycle. | Extended rotations, retention harvesting, selection harvesting | 65,983,843 | 36% |
| Even-aged management | Even-aged management comprises of a repetitive rotation cycle of distinct phases, including the regeneration, intermediate treatments (incl. thinning) and final harvesting. | Clearcut, clearcutting, even-aged management, even-aged stands, seed tree removal, two-aged management, extended rotations | 29,521,224 | 16% |
| Retention | Harvesting method in which some structural elements are retained at the time of harvest, such as mature trees and dead wood to increase the structural complexity of the stand. | Basal area and diameter retention, canopy retention, greater retention, overstory retention, retain biomass, retain dead wood, retain dead wood and recruitment trees, retain dominant and co-dominant trees, retain harvestable stock, retain recruitment trees, retention harvesting to promote shade-intolerant species, retention of wildlife and recruitment trees, single tree retention, variable retention | 87,817,348 | 47% |
| Selection harvest | Individual trees or smaller groups of trees are removed instead of all trees. Produces stands with several age-classes. | Group selection, hardwood control, hardwood release, improving species composition, increase standing and lying dead wood, old growth protection, release of well stocked conifer stands, rotation harvesting limited by basal area, selection cuts, selection for hardwoods and loblolly pine, single-tree selection, transition harvest | 38,008,407 | 21% |
| Regeneration | Re-establishment of the forest stand, through natural (from existing seeds, samplings in the stand) or artificial regeneration (planting, direct seeding) | Natural regeneration, planted seedlings, planting, prescribed burns, reforestation/replanting, regeneration harvesting with reserves, rehabilitation, rehabilitation of understocked areas, replanting shelterwood, shelterwood regeneration, shelterwood system | 17,777,969 | 10% |
| Pre-commercial thinning | Removal of specific trees or age-class of trees before trees reach merchantable size. | Pre-commercial thinning | 2,245,671 | 1% |
| Other thinning | Removal of specific trees or age-class of trees to improve the growth or health of the remaining trees. | Commercial thinning, intermediate thinning, variable-density thinning | 17,240,143 | 9% |

Forests are also facing a growing number of stressors that threaten C stocks ^12,13^. As the climate changes, these stressors pose increasing challenges for meeting the other requirement of an offset: *permanence*, or the persistence and longevity of stored C from offset projects. In addition to changing temperature and precipitation patterns, mega-disturbances, such as fires and insect outbreaks, are increasing in intensity and frequency ^13–15^. For example, in California, USA, eight of ten largest fires on record occurred within the past five years, burning almost seven million acres and releasing an unprecedented amount of CO_2_ into the atmosphere ^16^. Recent fires such as these raise concerns about the permanence associated with the forest carbon offsets, including where and how many forest carbon offsets could be reversed due to fires and other disturbances.

Forest management could play a role in reducing or exacerbating risks of carbon offset reversal from wildfire. Indeed, improved forest management practices differ in their ability to reduce risk of carbon loss from fires. For example, some practices, like extending harvest rotations, have been featured as a NCS pathway but may exacerbate risk of future carbon losses by retaining higher densities of aboveground biomass – some potential fuel for wildfires – longer in the landscape. In contrast, other practices like thinning reduces fuel loads by removing flammable biomass from forested landscapes. Yet, we lack a comprehensive understanding of risk to current offset projects over large spatial scales. Thus, the type and location of forest management practices will determine a project’s vulnerability to C losses and offset reversal.

The types of forest management practices being implemented are likely to determine the additionality and permeance of offsets, creating a pressing need to evaluate which forest management practices are being credited for offsets. In response, we compile a new database of forest management offsets from the only offset market in the U.S. to address: 1) which type of forest management is applied in existing forest carbon projects, 2) what is the ownership structure in these projects, and finally, 3) what proportion of offsets are at high or moderate risk from wildfire? For our final question, we focus on threats to current offset permanence from wildfire, rather than other disturbances, because it is the major disturbance on forestland in the Western U.S. where many forest carbon offset projects are located. Wildfires are also expected to increase in extent, intensity, and frequency as a result of climate change, thus threatening forestland across the US ^17–22^. Our analysis advances understanding of trends in forest carbon offset projects in the US by offering new details and perspectives for the of forestry projects involved in the offset credit market and assess the potential for carbon losses stemming from projects in wildfire prone areas.

## RESULTS

As of November 2020, 92% of issued offsets issued by California Air Resources Board, originate from forest carbon offset projects ^8^. Furthermore, 96% of these forestry projects are considered improved forest management and while, avoided conversion forest projects account for just 4% of the offsets issued. Improved forest management projects are heavily concentrated in the Western U.S.: 58% of forest offsets issued are from projects located in Alaska (AK), California (CA), Washington, Arizona, and Oregon, with the AK and CA accounting for 40% of issued forest offset credits (Figure 1, Table S1).

**Figure 1.**
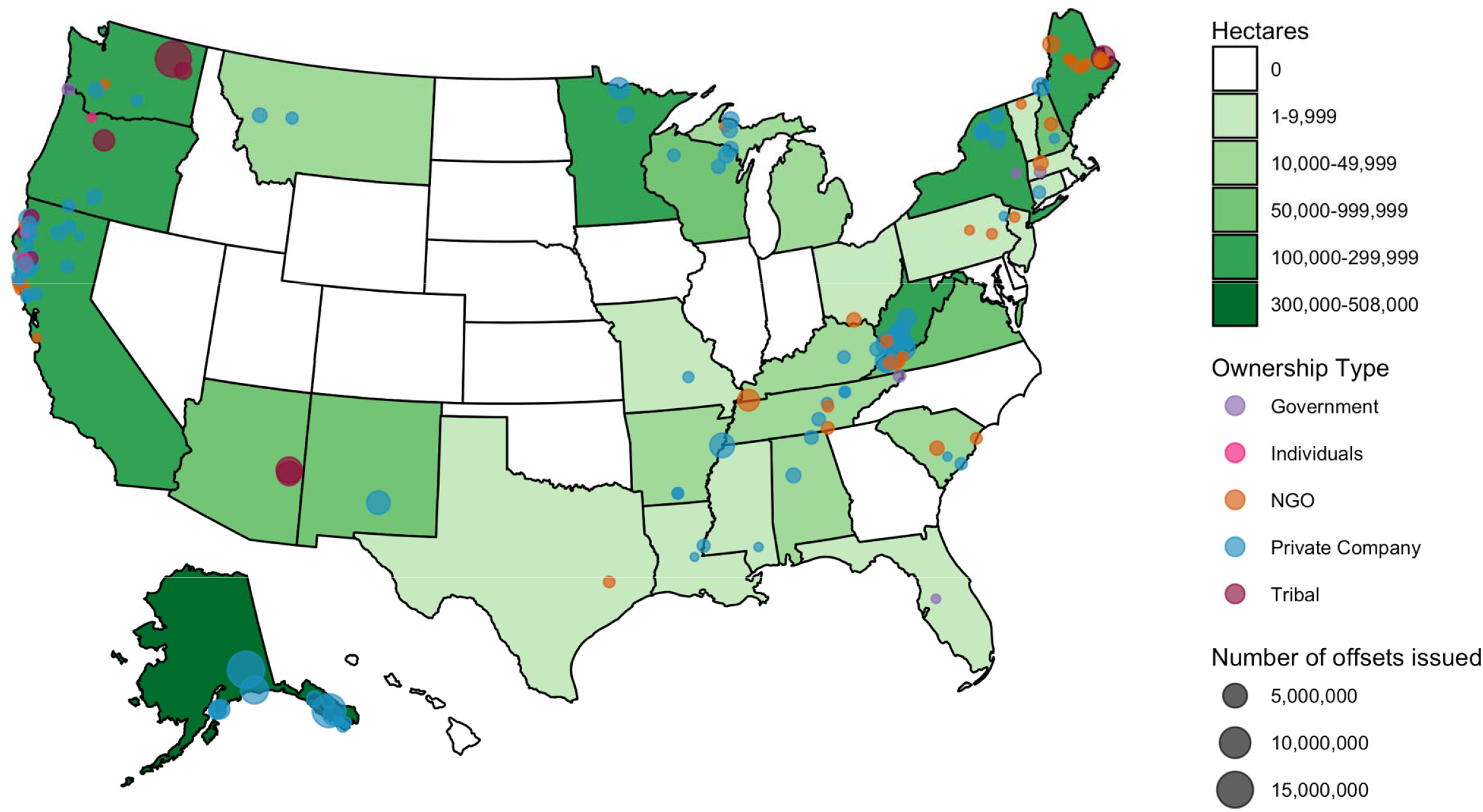
Locations of existing forest carbon offset projects (green, in hectares) in the United States and per ownership group (dots).

### Trends in Improved Forest Management offset projects

Improved forest management projects received a total of 185,088,866 credits, corresponding to 185 million metric tonnes of CO2. Improved forest management projects represent 96% of all forestry sector credits and 58% of all credits issued by the two offset project registries (OPRs) (Table S1). We identified 257 projects listed as improved forest management projects with American Carbon Registry or Climate Action Reserve, covering a total of 8,442,750 acres. Of those projects, 165 had been issued offset credits as of late 2020, and these existing or past projects covered 5,778,774 acres. Individual projects acreage ranged from 117 to 506,729 acres. Offset credits awarded to individual projects ranged from 2,616 to 15,456,787. Offset credits issued per acre ranged from 1.6 to 274, with a forest regeneration project in Mississippi receiving 808 credits per acre. Three projects were located outside of the United States, with one in Brazil and two in Madagascar. All other projects were distributed across 31 states (Figure 1, Table S1).

### Forest Management Practices

Almost half of all projects mentioned using retention harvesting, whereas 34% of projects listed no management or no commercial management, 36% listed uneven-aged forest management practices in their project documentation, and 16% of projects used even-aged management practices (Table 1). Selection harvesting was listed in 21%, precommercial thinning in 1%, and regeneration in 10% of projects, respectively, and 9% mentioned another thinning practice. Many projects listed multiple management strategies and therefore are counted in multiple categories for forest management.

Projects using no management or no commercial management of land accounted for 62,721,277 offset credits and 1,712,089 acres (Table 1, Tables S1 and S2). Not all projects mentioned previous land use or history, but many had been managed for timber harvest. On average, these projects without management or commercial forest management received the most credits per acre – 46 credits per acre. Even-aged management was applied in 955,323 acres and these projects received in total 29,521,224 offset credits, averaging 31 credits per acre. Projects mentioning uneven-aged management were listed for projects covering 1,998,772 acres receiving an average of 33 credits per acre with 1,056,534 acres managed, at least in part, by selection harvesting, with these projects receiving 36 credits per acre. Some of form of retention harvesting was practiced in 3,368,514 acres, receiving on average 26 credits per acre – it is important to note that retention harvesting was listed as a management practice in even-aged and uneven-aged forest management projects. Projects using pre-commercial thinning on 67,103 acres received 34 credit per acre on average, while those using other types of thinning received 16 credits per acre. Other type of thinning was used on projects covering 1,095,299 acres. Projects using regeneration practices totaled 1,056,040 acres and received about 17 credits per acre.

### Ownership

Companies own 75% of forest carbon offset project acres and received 69% of all credits issued (Figure 3, Table S2); four projects owned by Alaska Native Regional Corporations received 17% of all private company offsets and comprised 12% of all private company acres. These four projects were not managed for large-scale, commercial forest harvest. Native American tribes own comparatively few projects but own the next largest amount of project land (15% of acres) and received 21% of offset credits. Non-governmental organizations owned 9% of offsets and 10% of project acres. Government organizations or municipalities owned few projects and acres but received over 1 million credits (0.6% of acres and offsets). Less than 0.3% and 0.7 % of projects acres and offsets respectively were owned by individuals or universities combined. We found that a majority of the forest offsets are bought by private companies (Figure 2, Table S2), and these projects had no management or no commercial harvest in almost half the projects, with uneven-aged management was applied in third of the projects.

**Figure 2.**
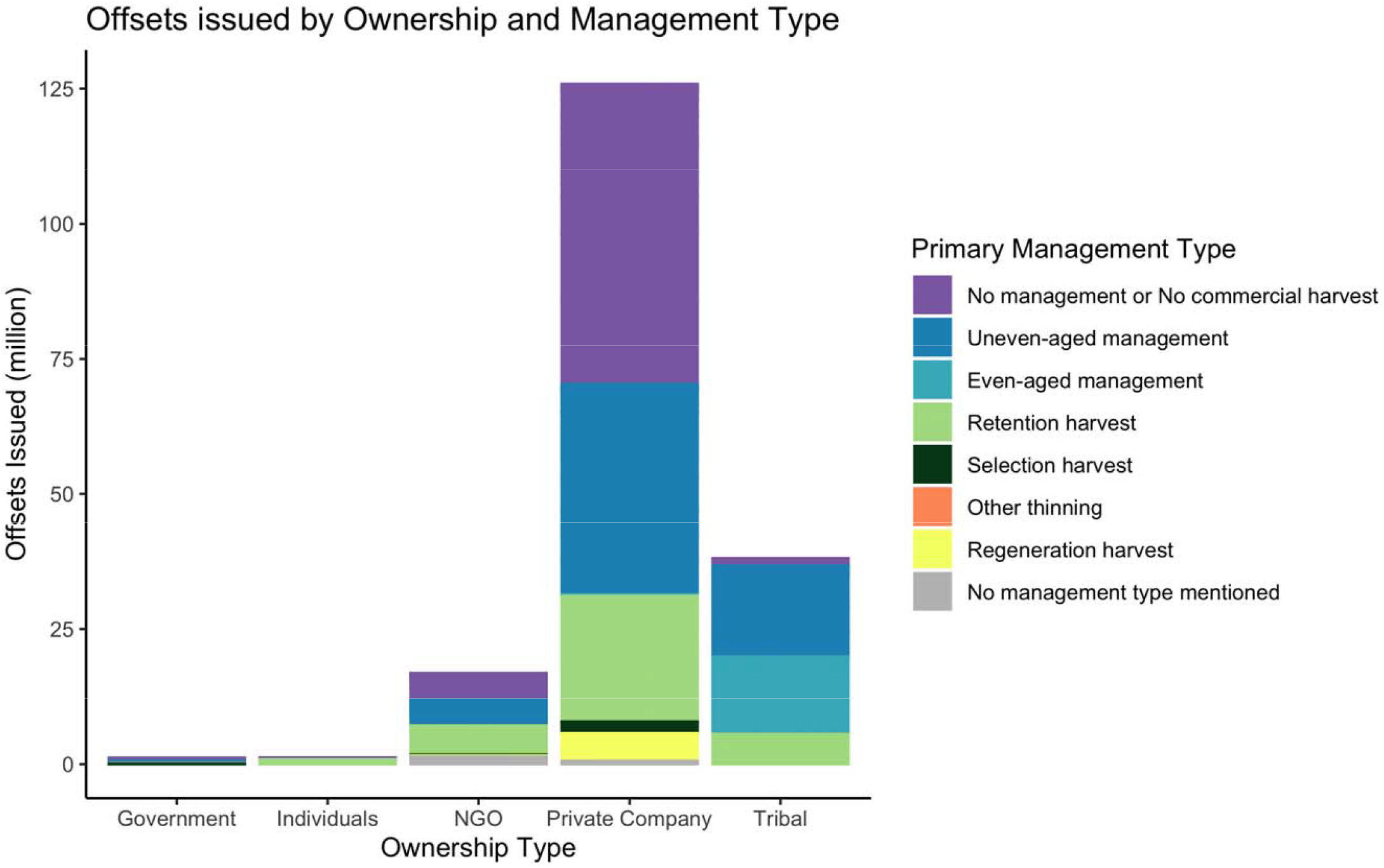
Forest carbon offsets issued per ownership group and forest management type. See Table 1 for descriptions of different types of forest management.

**Figure 3.**
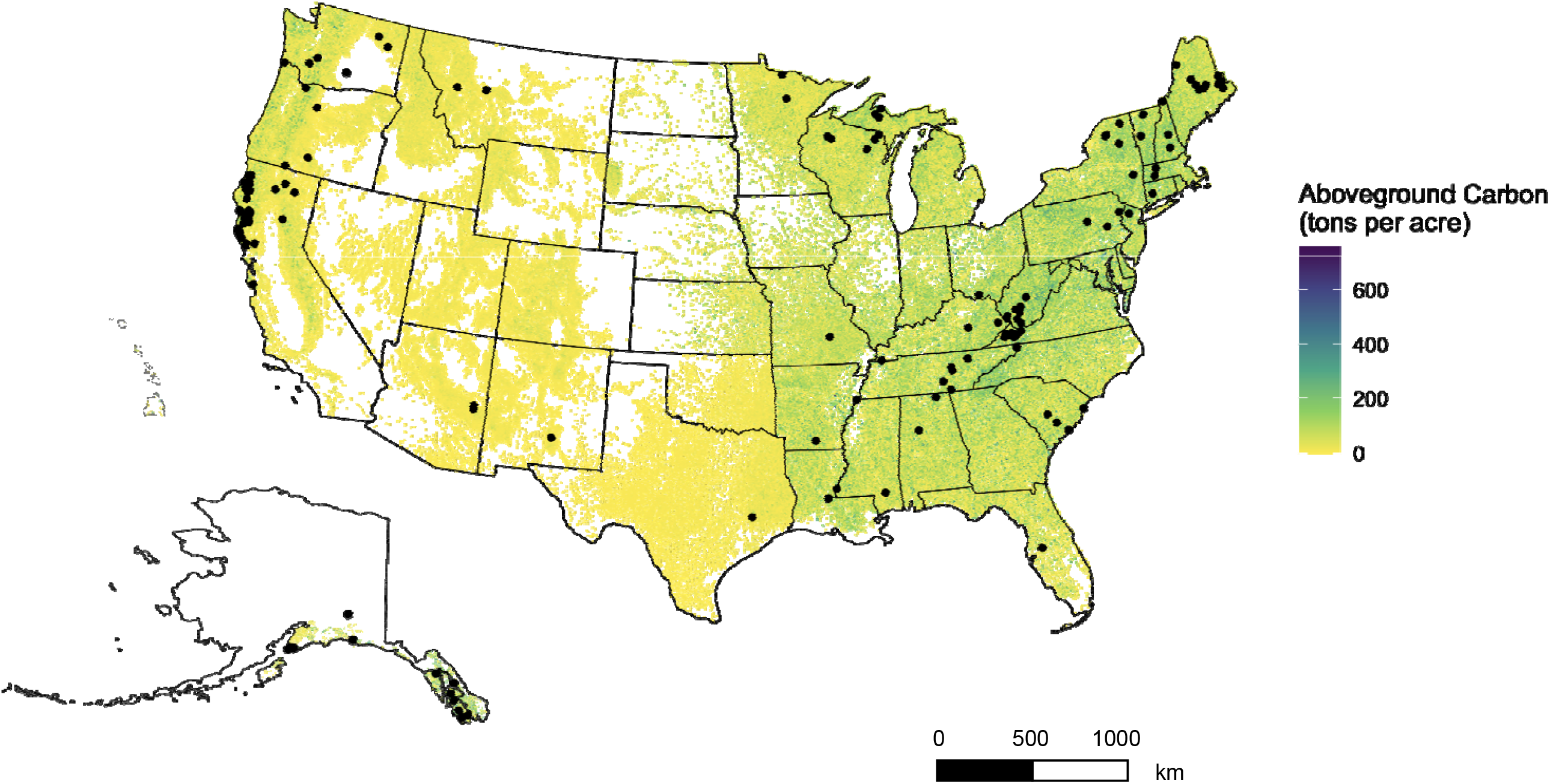
Map of Aboveground Carbon in the United States with locations of existing forest carbon offset projects. Aboveground C (US tons ac^-1^) (n = 11,674,137) in the United States and locations of (black dots). Data used for figure is from USDA Forest Services National Forest Inventory data (n = the number of samples).

### Risk of carbon losses from wildfire

In the U.S., 1,100,485 project acres – or 19 % of all forest project acres and 26% of forest project offset credits – are in areas of moderate wildfire risk accounting for 48,683,288 of issued offset credits (28% of all improved forest management offset credits) (Figure 5; *see Methods for more details*). Out of these projects, 46 projects – representing 16% of all forest offset credits and 9% of all forest project acres in the country – are in California. These projects account for all of project acres and project credits in the state. Other moderate risk projects are located in Oregon (1 project, 4% of project acres and 68% of credits in the state), South Carolina (1 project, 14% of project acres, 20% of credits) and Washington (2 projects, 98% of project acres and 95% of credits). Improved forest management projects tend to be located in areas with higher aboveground carbon densities, i.e., on forestland (Figure 2). Due to the productive nature of these forestlands, these project locations also tend to have high soil organic carbon and litter carbon densities (Figure 4).

**Figure 4.**
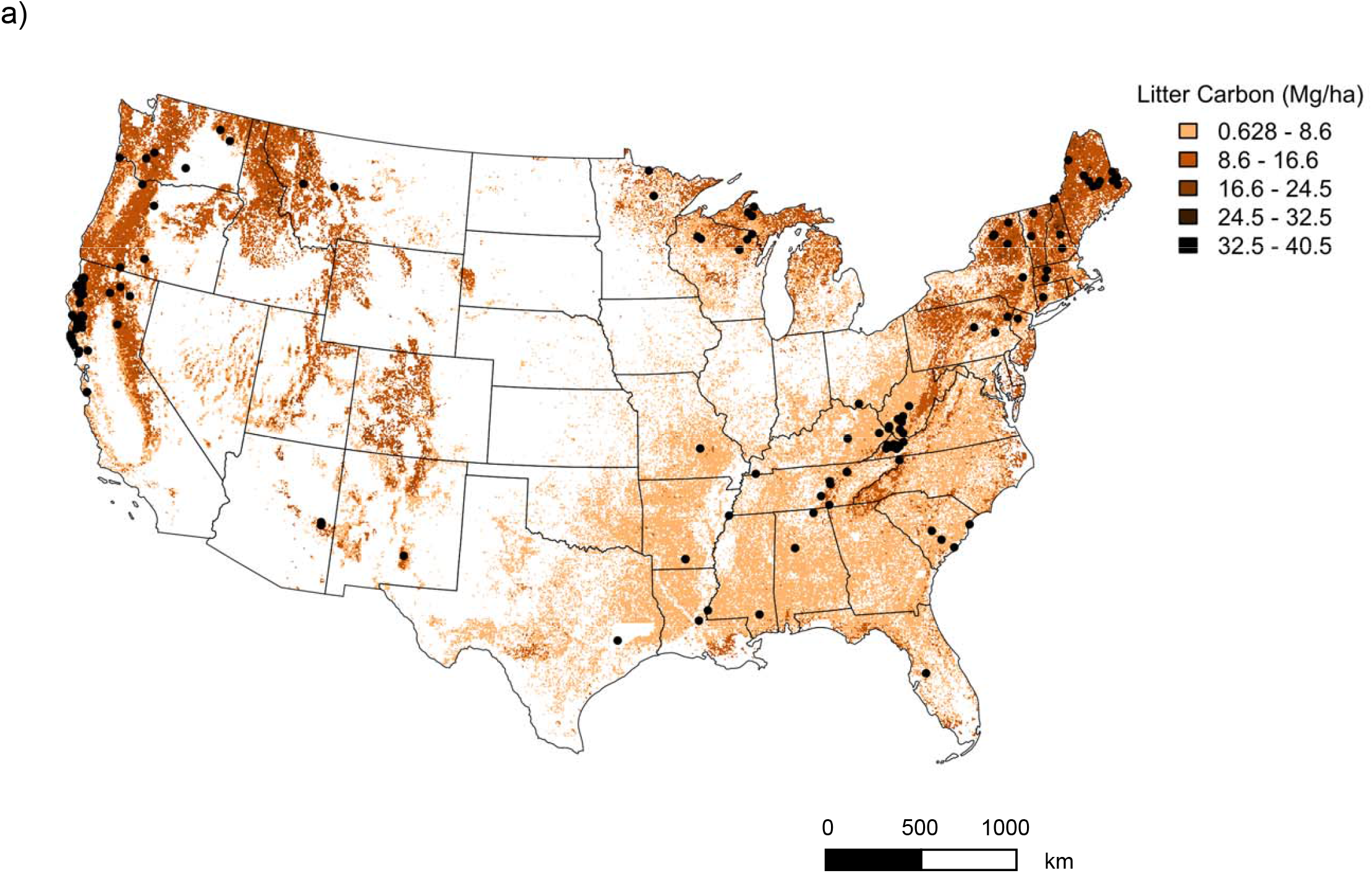
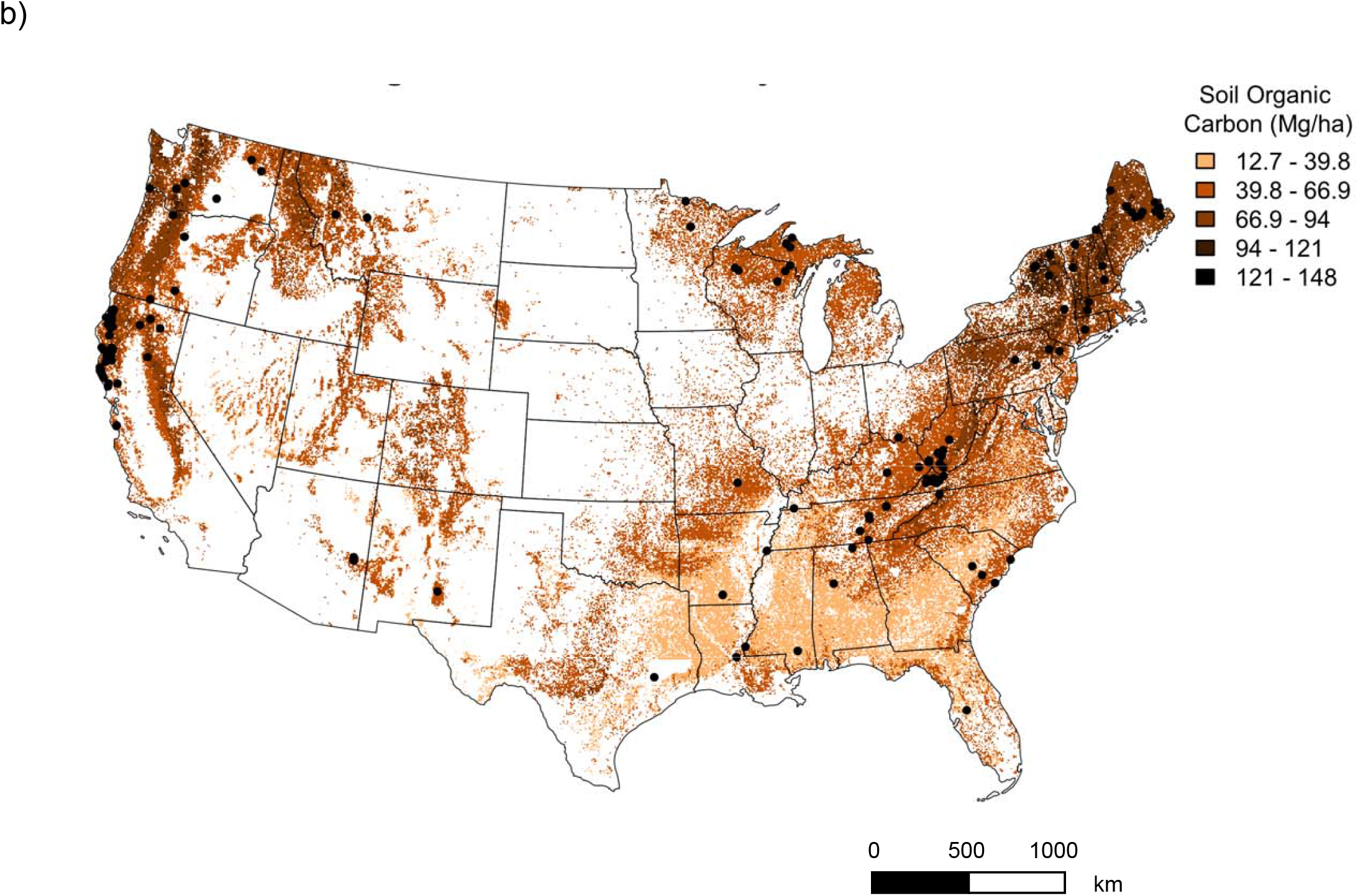
(a) Litter Organic Carbon (Mg ha^-1^) (n= 3303) and (b) Soil Organic Carbon (Megagrams ha^-1^) across continental United States and locations of existing forest carbon offset projects (black dots). Data used for figure is from USDA Forest Service National Forest Inventory data (n = the number of samples).

**Figure 5.**
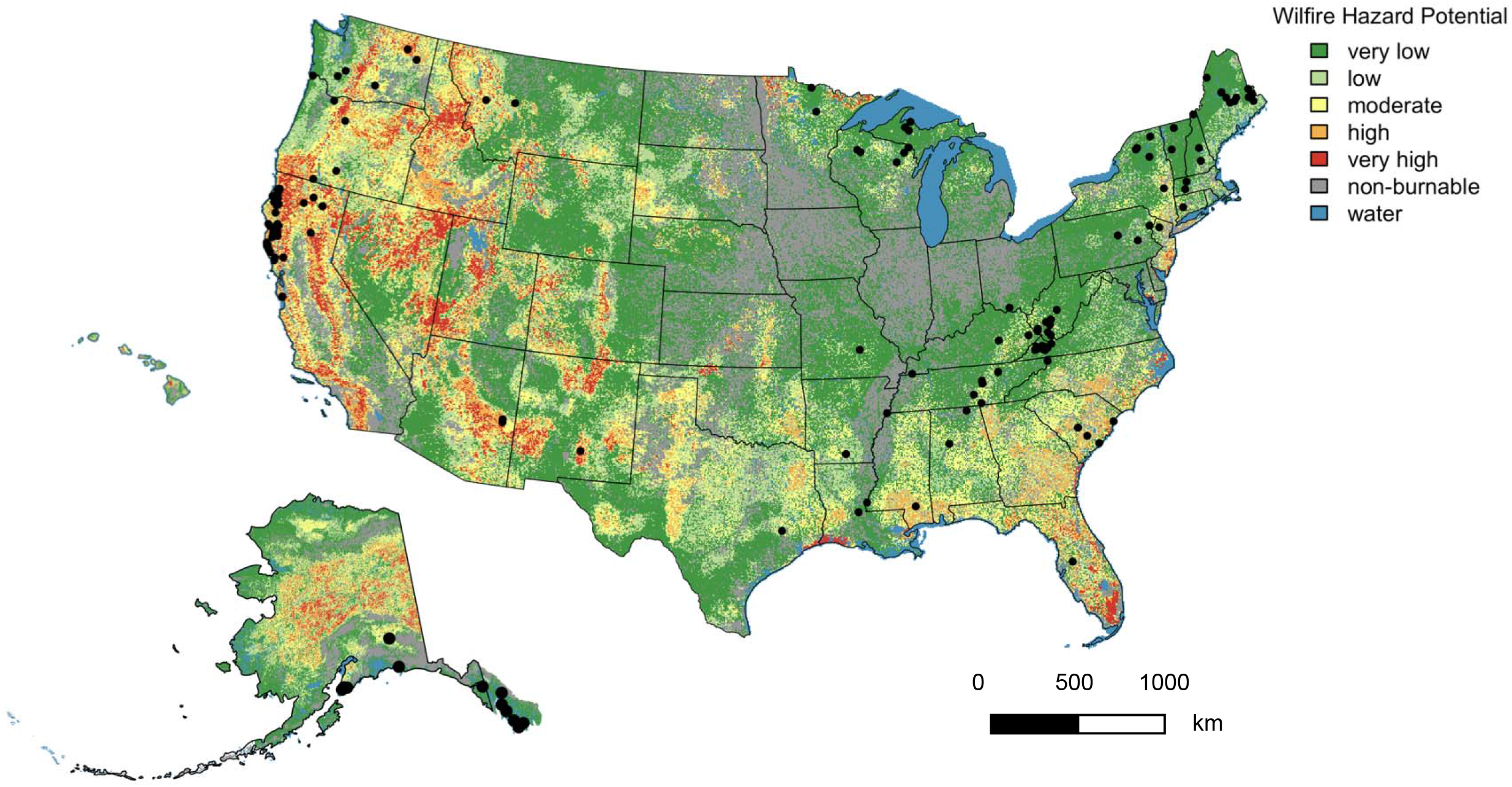
Wildfire hazard potential (WHP) in the United States and locations of existing forest carbon offsets projects (black dots) (WHP based on Dillon et al., 2020).

## DISCUSSION

Forests as a natural climate solution have dominated the discourse on climate-focused land management. Analyzing existing forest carbon offset projects in the US, we find that forest-based offsets are the dominant offset type in the US market, and improved forest management projects account for 96% of forest offset credits and 60% of all offset credit issued by the OPRs. We found that projects that list no management or no commercial harvest received the highest number of offsets per acre, followed by retention and selection harvesting. In addition, we observed that forest offset projects are indeed located forests with higher above- and belowground carbon densities (Figure 3 and Figure 4) but also areas of moderate and potentially increasing wildfire risk (Figure 5).

This analysis complement, buts differs from, recent research on natural-climate solutions from forest management ^2,11,23,24^. Prior analyses have primarily focused extended rotations as the NCS pathway from forest management ^2,11^. For example, Fargione et al. (2018) and Griscom et al. (2017) only include extending rotations in their analyses of NCS from forest management – yet those practices can increase exposure to risks of catastrophic carbon losses from disturbances and miss many other forest management practices currently being certified for offsets (Figure 5). Here, we broaden the view on forest management practices that could be considered effective and sustainable in managing forest carbon, based on practices being certified on-the-ground in carbon offset markets (Table 1). This expanded view is important because 96% of all offsets are from improved forest management projects, and these projects include a diverse suite of practices (Figure 2). Most projects do not mention if extended rotations are happening as part of ‘no management or commercial harvest’ (Figure 2). From our broader analysis of improved forest management implemented on the ground, we also highlight several gaps and research needs for offsets and natural climate solutions from forest management discussed next.

### Risk of reversal due to wildfire

Our analysis reveals that 28% of improved forest management projects are areas with moderate fire risk (Figure 5). While our analysis did not find any existing projects with very high or high wildfire hazard potential, we did find a project in California located on forestland with high wildfire hazard potential, which had been issued 847,985 offset credits, but the project was terminated in 2019 due to a wildfire. Yet, we anticipate increasing risk to offsets from fire and other disturbances, which are increasing in frequency and intensity with climate change ^12,13,25^. For wildfire specifically, the convergence of warming temperatures and expanded ignition pressure from people is increasing the number of large human-caused wildfires and the fire-niche across the Western US ^26^ (Figure 5). As a result, wildfires could threaten carbon offsets from forests across the U.S. – not just in the flammable West. Increasing demand for forest based offset credits could also drive the expansion of projects further into fire-prone landscapes, where fuel conditions are further exacerbated by the unpresented drought ^27–29^. Finally, predictions of future wildfire occurrence and outcomes are inherently uncertain ^30^, adding to the uncertainty associated with forest carbon offset permanence.

While low- and mixed-severity fires have historically been a natural phenomenon in the forested ecosystems of the Western US, human influence (e.g., grazing, land conversion, urbanization, fire suppression) has resulted in exclusion of fires in the region ^31,32^. Decades of fire suppression have altered the structure and composition of many forests in the western United States, some of which are now also facing the compound disturbance effects of fire, bark beetles and drought ^27^. In fire-suppressed forest stands, uncontrolled, high intensity wildfires tend be severe in terms of their impact on aboveground carbon stocks. Forest stands with high aboveground carbon densities tend to be more vulnerable to forest fires due to overstocking of flammable biomass following fire suppression. If a high-intensity fire were to occur, carbon losses from these stands could be significant. if left untreated for fuel, forestland can release more CO_2_ once they burn than thinned ones as large, as catastrophic wildfires tend to consume all available biomass, including the litter layer and surface layers of the soil ^32,33^.

These findings about CO_2_ emissions from untreated (i.e., no fuel management) forestland with legacy of fire suppression have important implications for many forestry projects in the offset credit program. First, forest carbon offset projects are situated within areas of high above- and belowground carbon densities (Figure 3, Figure 4), suggesting that these areas (i.e., forest stands) have been excluded from fire or other large-scale disturbances in the recent past. Second, many locations had been previously managed for timber harvest thus implying that the initial or project start aboveground carbon stocks were considerable. Finally, we find that a disproportionally large number of forest carbon offset project land (i.e., forestland) is left untreated or not managed, and just a few explicitly use thinning or other types of fuel management practices.

Guidelines for managing fuels on these forestlands participating in offset programs is urgently needed given the risks – particularly in California and across the western states where wildfires are common. To date, only six existing forest offset projects in our database mention prescribed burning as a management practice and only one of these projects was in the West (in New Mexico). To that end, we recommend that improved forest management further expand its definition (reviewed in ^7^ to include active fire and fuel management. In addition, future markets from other sectors (e.g., agricultural crop losses) that face losses from events like drought could provide some guidance for these emerging markets ^e.g., 34^.

### Finding a sustainable path for forest carbon offsets

In operational forestry, improved forest management is not well-defined, and the long-term carbon benefits of most forestry practices considered improved forest management remain to be tested ^7,9^. Currently, markets are certifying forest offsets projects but offer limited accountability and transparency for additionality – the demonstrated effects of carbon sequestration in the forest stand under improved forest management practices. Large quantities of offset credits are awarded to projects at the start of the project (i.e., initial tracking period), particularly for improved forest management projects ^35,36^. Currently, there are no policy instruments or regulation in the California offset credit market focusing on oversight and accountability of forest offset projects, and the governance for environmental integrity is focused on the development (i.e., the protocols) and start of the forest project ^36,37^. The current process has put into question the added carbon benefits of these projects ^36^

Our analysis reveals that credited projects vary to a great degree in their disclosure about the planned or completed forest management activities for the project area. While several forest carbon offset projects provided detailed descriptions of the management objectives, and by that extension forest management practices, many offered little detailed information on what type of management activities will take place and when. For example, when managing for forest carbon, there is considerable ambiguity associated with practices listed “retention” (Table 1). However, *what* is retained in the forest stand and for what *purpose* was not implicitly highlighted in the forest carbon offset project documentation. Going forward, a thorough and transparent planning and monitoring network for the forest practices applied in these projects, including retention harvest practices, would aid in determining the extent and scale of additional carbon benefits. Forming a new partnership between the entities involved with the carbon offset market, including the state of California, developers of the forest offset protocols (OPRs), and finally, the forest research community could assist in building a framework for assessing forest carbon offset opportunities on forestland in California.

Finally, we recommend several future directions for research, partnerships, and policies around forest management offsets. While assessing the effectiveness of California’s offset market to demonstrate significant carbon emission reductions is beyond the scope of this article, we call for substantial investments into oversight of credited and existing, and future forest offset projects. Our results further highlight that a significant portion of existing forest carbon offsets face a risk of reversal through wildfire, including all the existing projects in California. Future research and partnerships could build a body of evidence for not only how these improved forest management strategies impact carbon, but also the extent to which they mitigate or exacerbate risk from disturbances such as wildfire ^31,38,39^ and pest outbreaks ^29^. For example, some improved forest management strategies such as thinning reduce fuel loads, but thinning only represented around 10% of credits, whereas no management of projects could increase exposure to risk of catastrophic carbon losses and was the dominant practice in 34% of certified projects (Table 1). Specifically including climate-driven disturbance risks such as wildfire in the forest offset protocol could increase the robustness of the offset credit program and help accurately determine the risk associated with each project. From a policy perspective, these results underscore that more sophisticated insurance mechanisms are needed for forest carbon offset losses and reversals, as well for the validation of long-term carbon benefits from different types of forest management.

## MATERIALS AND METHODS

We review trends in forest carbon offsets, in terms of the types of forestry management practices listed, land ownership, number and location of issued offset credits, and potential risk to these offsets from wildfire. To do so, we compiled a novel database of all forestry projects from the two largest offset project registries in the US’s only carbon market (California’s Cap-and-Trade and Voluntary Offset programs): the Climate Action Reserve (CAR) and the American Carbon Registry (ACR). In these programs, an offset credit represents an emission reduction of one metric tonne of CO_2_.

### Offset Databases

As of February 2022, 214,981,710 forest carbon offset credits in total (totaling to 215 Tg CO_2_e) have been issued through California Air Resources Board ^8^. In November 2020, we accessed, downloaded, and compiled the data from two largest offset project registries (OPRs) in the United States; the Climate Action Reserve (CAR) and the American Carbon Registry (ACR), which track offset projects and issue offset credits, and are responsible for verifying and certifying emission reductions (see SM for details). We examined all “Forest Carbon’’ projects from the CAR, and all “Improved Forest Management” (IFM) projects from the ACR. Our database included 143 Forest Carbon projects from CAR (as of November 16, 2020) and 139 IFM ACR (as of December 4, 2020). Information on the project ID, developer, name, owner, year registered and/or listed, status, ARB status, site location and number of offsets issued where retrieved directly from the OPRs.

From each project document we gathered information on management practices and project acreage (Table 1; for more details, see Supplementary Materials). Practices were sorted into 8 main categories based on the definitions from the Southwest Fire Science Consortium’s silviculture terminology and the US Forest Services’ reforestation glossary (see Table 1). Ownership type was assigned based on information in the project submittal form. If more information was required, the organization’s website was referenced. Project documents were reviewed in November and December 2020. Management information was not available for 32 projects. Of those, 4 had been canceled, 4 were inactive and 2 were completed. Three projects with no management information had received offset credits. We contacted project owners for 25 projects and heard back from owners of 21 projects but were unable to attain additional information on management practices. Our analysis includes active offset projects that have received offset credits and includes completed projects but not planned (ARB-status listed as *proposed*) or inactive projects (ARB-status listed as *inactive*). We obtained project coordinates from project paperwork and approximated coordinates when not available (see SM for details). From this data, we calculated the total credits and acreage in each state and in each management and ownership categories.

### Litter and soil carbon maps

We overlaid project locations on maps of soil carbon and litter carbon data from Cao et. al. ^40^ and onto maps of aboveground carbon calculated from the USDA Forest Services’ National Forest Inventory data ^41^ and the ‘rFIA’ package v3.1 ^42^ (see SM for details). All analysis and figures were completed using R version 4.0.4 (R Core Team, 2021).

### Wildfire risk for existing forest projects

We overlaid forest offset project location data with Wildfire hazard potential (WHP), retrieved (February 15, 2021) from USDA ^43,44^. Dillon and Gilbertson-Day ^43^ modeled WHP using spatial datasets of wildfire likelihood and intensity generated in the Large Fire Simulator program, spatial information on fuels and vegetation data from LANDFIRE 2014 and point locations of past fires. Each project’s county mean categorical WHP was used as a proxy for the project area WHP. For projects in multiple counties, the county WHPs were averaged and each WHP-value, if a fraction, was rounded down. State WHP was used for projects that did not specify a county or that were in multiple states. Based on this WHP data, greater than 4 on a scale of 0–5 is considered a high fire risk location, whereas 3 is considered a moderate risk location.

## Funding

The authors thank the Finnish Cultural Foundation for funding to LK

University of Colorado Boulder Undergraduate Research Opportunity Program for supporting JR.

## Data and materials availability

Data will be published as supplementary files upon acceptance of the manuscript and was made available for reviewers to evaluate the manuscript. Code to produce the figures can be found at the project GitHub page: lillikaarakka/ifm_projects_2022.

