## Supplemental Materials for "Trends in forest carbon offset markets in United States"

**Methods: Additional Details on Review**

*Public Registries for offsets*

Climate Action Reserve, 2020. Public Registry of Offset Projects. <https://thereserve2.apx.com/mymodule/mypage.asp> (Accessed and downloaded December 4, 2020)

American Carbon Registry, 2020. Public Registry of Offset Projects. <https://acr2.apx.com/myModule/rpt/myrpt.asp?r=111> (Accessed and downloaded November 16, 2020)

*Project details*

We examined and reviewed mentions of management practices and project acreage in the project design document or project plan, if available, or the project submittal form. Management practices were most often found in the project overview, project eligibility, or similar section in project design or project plan documents. On project submittal forms for both OPRs, management practices were most often listed in Part V, section H, “describe the historical land uses, current zoning, and projected land use within the Project Area and surrounding areas,” or Part VI, section C, “Describe the management activities that will lead to increased carbon stocks in the Project Area, compared to the baseline.” All practices listed in the project documents were added to our database. In the body text of the article, companies are synonymous to *buyer*s [of carbon offset credits].

*Location coordinates*

For projects that did not list coordinates and for two projects (ARC192 and CAR1195) whose listed coordinates were in the ocean, approximate project coordinates were obtained. The project location, generally a city or county, listed on the project submittal form was uploaded to Google Earth and the associated latitude and longitude exported. For locations Google Earth could not parse, for example when a project listed miles away from a town in a specified direction, Google maps was used to approximate the location for mapping purposes.

*Aboveground carbon calculations*

Aboveground carbon data was calculated from the Plot and Tree tables from the US Forest Services’ Forest Inventory Analysis (FIA) dataset (Gray et al., 2012). The r package rFIA version 0.3.1 was used to calculate aboveground carbon for FIA plot in each state, using the protocol suggested on the rFIA website (Stanke et al., 2020). The average aboveground carbon per tree in pounds was multiplied by trees per acre and divided by 2000 to result in tons of aboveground carbon per acre.

*Project documents*

Our analysis reveals that while improved forest management is the most credited project type in the CA market, existing projects vary to a great degree in their disclosure about the planned or completed forest management activities for the project area. While several forest C offset projects provided detailed descriptions of the management objectives, and by that extension forest management practices, for the project area, many offered little detailed information on what type of management activities will take place and when. This is an interesting contrast to California’s existing environmental protection regulation associated with forest harvesting, including environmental impact assessment requirements such as the timber harvesting plans (mandated for any type harvesting on private lands in California by state’s Forest Practice Act), which are prepared by licensed forestry professionals. Not all projects mentioned previous land use, but many had been managed for timber harvest thus implying that the initial or project start aboveground C stocks were considerable.

*Analyses and visualizations*

Trinity Timberlands University Hill Project (CAR1046) was omitted from analysis, as it was listed as terminated in the offset database. Finally, Boa Vista project (ACR105) was removed from analysis as the project was outside of US. Bishop Improved Forest Management project (CAR973 – 3,112,189 offset issued between 2014–2020) was not included in the analysis as we were unable to verify the acreages and management practices for the project, and did not receive a response from the project developer upon contact.

FIA-database version 8.0 (downloaded 5/27/21) was used to estimate litter and soil carbon stocks. All visualizations were done in R version 4.0.4 using the packages ‘usmap’ version 0.5.2, ‘ggplot2’ version 3.3.5 and ‘terra’ version 1.3.4 (R Core Team, 2021).

**Calculation of risk [of reversal] for offset projects**

Offset project registries risk assessment for the determination of a forest project’s reversal risk. Permanence and risk of C loss factor into the credits as a risk rating percentage (varies between 2–8% depending on the OPR), which is used to determine how many offsets the project must contribute to the buffer pool. For example, ACR applies a 4% risk in areas with high fire risk and 8% if the project is in an area (<30 miles) where a larger fire (1000 acres) fire has occurred in the last 12 months. Similarly, CAR assesses project wildfire risk based on project location, but reduces the risk rating for projects that carry out vegetation treatments aimed at reducing fuel loads. Like fuel treatments, all forest management activities require an investment from the forest owner.

**Climate action reserve** – Risk Rating Analysis (2019):

The project’s reversal risk rating is calculated as follows:

Risk of Reversal % =

10- − [(1 – Financial Failure %) × (1– Illegal Forest Biomass Removal %)

× (1– Conversion %) × (1 – Overharvesting %) × (1 – Social Risk %) × (1–Wildfire/Disease/Insect Outbreak %) × (1–other Catastrophic Events %)]

**American Carbon Registry** – Tool for Risk Analysis and Buffer Determination (2020)

The project’s reversal risk rating is calculated as follows:

Total Risk score % =

Financial Risk + Project Management Risk + Social/Policy Risk + Conservation Easement Deduction + Natural Disaster Risk (area-specific)

**Values per category of risk:**

**Management and Governance Risk**

Financial – 4% default value, 3% US public and Tribal lands

Project Management – 4% default value, 3% US public and Tribal lands

Social/Policy – 2% default value, 5% if project is located outside of the US, 3% if project is located outside of the US and demonstrates community engagements through ACR-approved mechanism

Conservation Easement Deduction – -2% Default value, -3% if there is regular onsite monitoring of activities related to carbon-specific conservation activities

**Natural Disaster Risk**

Fire – 8% if the project is located in an area where fire greater than 1000 acres has occurred within 30-mile radius of project area in prior 12 months, 4% if project is located in high fire risk region, 2% if project is located in low fire risk region (verifiable evidence must be provided), 1% for agriculture and grassland projects only

Diseases and Pests – 8% if epidemic disease or infestation is present within project area, or within 30 mile radius of project area, 4% default value

Levee Failure and Water Table Changes – 2% Default for all wetland projects (and for forest projects where more than 6- of the project area is a forested wetland)

Other Natural Disaster Events – 2% Default Value for all sequestration projects

Table S1 – Forest Carbon Offsets issued per U.S. State (as of December 2020).

| **State** | **Projects** | **Offset credits issued** | **Offset project acres** |
| --- | --- | --- | --- |
| Alaska | 12 | 45,004,121 | 935,384 |
| Oregon | 5 | 4,762,028 | 663,209 |
| West Virginia | 8 | 22,918,984 | 574,047 |
| Washington | 4 | 16,815,558 | 559,025 |
| California | 46 | 29,321,219 | 522,997 |
| Maine | 13 | 8,262,760 | 478,079 |
| New York | 6 | 4,925,885 | 459,295 |
| Minnesota | 2 | 4,503,337 | 340,560 |
| New Mexico | 1 | 4,455,664 | 228,376 |
| Virginia | 9 | 5,436,414 | 170,288 |
| New Hampshire | 3 | 2,440,537 | 152,236 |
| Wisconsin | 5 | 3,501,665 | 134,822 |
| Arizona | 2 | 10,917,224 | 131,904 |
| Tennessee | 8 | 5,448,087 | 78,766 |
| Michigan* | 3 | 2,498,888* | 69,616 |
| Kentucky | 2 | 1,035,335 | 46,283 |
| Alabama | 2 | 1,452,678 | 40,731 |
| Arkansas | 3 | 5,495,898 | 37,918 |
| South Carolina | 4 | 1,296,983 | 32,690 |
| Montana | 2 | 1,074,933 | 31,815 |
| Massachusetts | 2 | 1,099,016 | 23,251 |
| Ohio | 1 | 684,708 | 15,474 |
| Mississippi | 2 | 460,353 | 9,913 |
| Vermont | 2 | 127,869 | 8,160 |
| Pennsylvania | 3 | 227,185 | 7,480 |
| Connecticut | 1 | 381,992 | 6,084 |
| Florida | 1 | 34,662 | 5,242 |
| Texas | 1 | 218,667 | 5,033 |
| Missouri | 1 | 170,586 | 3,982 |
| New jersey | 1 | 100,696 | 3,174 |
| Louisiana | 1 | 14,934 | 2,941 |
| **Total** | **156** | **185,088,866** | **5,778,774** |

*Project CAR973 in Michigan that had received 3,112,189 credits as of 2020, was excluded from the analysis, as we were unable to confirm the acreage of the project through project documentation and unsuccessful enquiries to the project developer, Blue Source LLC.

Table S2 – Forest Carbon Offsets issued per ownership group and forest management listed for the offset project (as of November 2020).

|  | No management / or no commercial harvest | Even-aged management | Uneven-aged management | Retention | Selection | Other thinning (not precommercial thinning) | Regeneration | No management mentioned | Project acres | Total offsets issued |
| --- | --- | --- | --- | --- | --- | --- | --- | --- | --- | --- |
| Private Company | 44% | - | 31% | 18% | 2% | - | 4% | - | 4,139,509 | 125,872,901 |
| Tribal | 4% | 38% | 44% | 15% | - | - | - | - | 1,280,960 | 38,136,101 |
| NGO | 29% | - | 28% | 32% | 1% | - | 1% | 8% | 561,469 | 16,862,056 |
| Individuals | 28% | - | - | 72% | - | - | - | - | 14,867 | 1,229,127 |
| Government | 50% | - | 35% | 8% | 8% | - | - | - | 35,899 | 1,185,259 |
| University | 100% | - | - | - | - | - | - | - | 2,673 | 78,069 |
|  |  |  |  |  |  |  |  | **Total** | 6,035,377 | 183,363,513 |

If a project mentioned several practices and to avoid double counting management categories for the table were ordered: no management/no commercial harvest > uneven-aged > even-aged> retention > precommercial thinning > selection > other thinning > regeneration > no management types.
